# Changing pollination rates affect plant life history strategies

**DOI:** 10.64898/2026.02.11.705202

**Authors:** Dylan T Simpson, William K. Petry, Paul J. CaraDonna, Amy M. Iler

## Abstract

An organism’s life history strategy is an attempt to optimize fitness, given environmental constraints and inherent demographic tradeoffs. As such, life history helps to shape an organism’s ecological and evolutionary responses to environmental change. However, life history can also be shaped *by* the environment, as the organism’s demographic rates respond—directly or through tradeoffs—to the new conditions. This feedback between life history and environment remains poorly understood, limiting our ability to predict the outcomes of environmental change. Here, we studied the effects of environmental change – specifically altered pollination services – on four perennial plant species. We conducted a field-based demography experiment that subjected naturally occurring populations of *Delphinium nuttallianum, Hydrophyllum fendleri, Potentilla pulcherrima* and *Erigeron speciosus* to three pollination treatments: ambient (control), reduced, or increased pollination. We estimated population growth rate (λ) and 11 metrics describing life history strategy and demographic resilience from an Integral Projection Model we constructed for each species and parameterized with 4–5 years of census data. Although most life history metrics responded idiosyncratically to pollination treatment, we found consistent effects of pollination on generation time, longevity and, in three of four species, recovery time. Specifically, reduced pollination led to increased longevity, generation time, and recovery time, and increased pollination led to the opposite. These changes in life history resemble shifts along the slow-fast continuum; reduced pollination led to slower lives and increased pollination led to faster lives. This is consequential because generation time and longevity influence short- and long-term population dynamics – for example, by affecting demographic stochasticity and sensitivity to environmental stochasticity, or rates of adaptation to novel conditions. Notably, these changes occurred largely independent from changes in population growth. Altogether, our results highlight changes in life history as an important but underappreciated consequence of environmental change.

## Introduction

An organism’s life history strategy is central to its ecology and evolution. Life history describes how and when an organism invests in growth, survival, and reproduction, which together shape population-level responses to the environment (Stearns 1976). For example, life history affects a population’s sensitivity to environmental variation (Morris *et al*. 2008) or particular environmental variables, (Paniw *et al*. 2018; Reznick & Endler 1982; Youngflesh *et al*. 2025), rate of recovery from disturbance, (Capdevila *et al*. 2020), and rate of adaptation to novel conditions (Martin & Palumbi 1993; Smith & Donoghue 2008; Vander Wal *et al*. 2013). Life history can also, however, be shaped *by* the environment (Reznick *et al*. 2019). Life history strategies are an organism’s attempt to maximize fitness, given environmental constraints and inherent demographic tradeoffs (Sæther *et al*. 2016). If the environment changes, so too might life history strategy, as vital rates respond—directly or through tradeoffs—to the new environmental conditions.

Study of life history evolution has found that life histories can respond rapidly to the environment through both adaptation and plasticity (Baird *et al*. 1987; Bårdsen *et al*. 2011; Felmy *et al*. 2022; Reznick *et al*. 2019). While these changes are not always predictable (Grainger & Levine 2022), environmental pressures that target specific life stages or demographic processes tend to induce life history tradeoffs. In a classic example from Trinidadian guppies, the presence of predators that target larger individuals lead to faster growth and earlier reproduction (Reznick *et al*. 1990). In frogs, elevated and more variable temperatures reduced juvenile survival, leading to delayed reproduction and greater investment in growth and adult survival (Reniers *et al*. 2015). Life history tradeoffs may thus play an important role in mediating responses to environmental change.

In contrast to studies of life history evolution, ecological studies of environmental change tend to focus on demographic vital rates or population growth (λ), rather than life history (e.g., Deutsch *et al*. 2008; Epps *et al*. 2004; Iler *et al*. 2019; McLaughlin *et al*. 2002; Radchuk *et al*. 2013; but see Williams et al. 2015, Cayuela et al. 2022, Burc et al. 2025). Changes in λ are certainly important – λ determines whether a population grows, shrinks, or remains stable – but environmental effects on life history could also have lasting impacts. For example, even with no effect on λ (e.g., Villellas et al. 2015, Cayuela et al. 2022, Blake and Coulson 2023), environmental change could affect aspects of life history like reproductive strategy (Baird *et al*. 1987; Kim & Donohue 2011), reproductive timing (Bernot *et al*. 2006; Reniers *et al*. 2015), or body size and longevity (Reniers *et al*. 2015). Longevity and reproductive timing determine generation time, which contributes to variance in λ and extinction risk (Morris *et al*. 2008; Pearson *et al*. 2014). Given the potential effects of the environment on life history, and its many consequences, ecological studies of environmental change would benefit from expanding their scope to include life history.

For plants, a key environmental determinant of fitness is pollination service by animals. Most plant species require animal pollination to reproduce (Ollerton *et al*. 2011), and plant reproduction is often pollen limited (Knight *et al*. 2005). Critically, many pollinator taxa are at risk or in decline (Cornelisse *et al*. 2025), which raises the question of how changing pollen receipt will affect plant populations. Changes in pollen receipt are often positively related to plant reproductive fitness components (e.g., Brosi and Briggs 2013, Lundgren et al. 2015, Thomson 2019), but the net effect on population-level average fitness, λ, can vary widely (e.g., Baer & Maron 2018; Knight 2004; Law *et al*. 2010; Iler et al 20XX). Seed production is resource-intensive, meaning that a change in the amount of pollen received could alter the relative investment in reproduction versus survival and growth (Van Noordwijk & De Jong 1986; Williams 1966). Even with no effect on λ, this reallocation could affect life history. For example, if plants invest less in reproduction due to reduced pollination services, individuals might live longer and slow the rate of turnover at the population-level. In a more extreme example, changes in rates of survival versus reproduction have led populations to shift between semelparity and iteroparity (Kim & Donohue 2011; Paige & Whitham 1987). More fully understanding the consequences of changing pollination services might therefore require looking beyond just vital rates to consider emergent life history traits.

Here, we use a multi-year demography experiment to examine the effects of pollination rate on life history strategies of four perennial plant species. We used data from this experiment to parameterize integral projection models, then use these models to estimate population growth rate (λ) and life history traits for each species under ambient, reduced, or increased pollination. We asked whether changes in pollination lead to changes in life history strategy and, if so, which life history traits are changing. Further, to determine whether any of these changes might be generalizable, we asked whether any changes in life history consistent are across species.

## Methods

### Overview

We estimated life history metrics and λ for four focal species under different experimental pollination treatments using Bayesian Integral Projection Models (Elderd & Miller 2016). Within each species, we tested for life history responses to pollination treatment using canonical discriminant analysis – a form of ordination – to summarize how life history metrics and λ change in multivariate space. We also directly compared the value of each metric in each treatment against the control. We then compared treatment effects among species to identify general patterns in life history response to altered pollination. The experimental design and Integral Projection Modeling are fully described in Iler et al. 20XX, and we briefly summarize these components below.

### Field-based demography experiment

To test for effects of changing pollination rates, we performed a field-based experiment using four perennial, subalpine herbs: *Delphinium nuttallianum, Erigeron speciosus, Hydrophyllum fendleri* and *Potentilla pulcherrima* (referred to hereafter by genus only). We chose these species because they differ in life history and dependence on pollination for seed set (Table S1). We performed the experiment at the Rocky Mountain Biological Laboratory (Gothic, CO, USA) from 2017-2022. Plots contained naturally occurring populations, which meant the number of plots per species varied to reach a target of 250 individuals per species and treatment; there were 16 plots for *Hydrophyllum*, four plots for *Potentilla*, and six plots each for *Delphinium* and *Erigeron*. Plots were 1-m x 10-m in size, arranged in a block design where plots in a block were directly adjacent and treatments were randomly assigned among plots.

The experiment followed individual plants, each tagged with a unique ID number, in three treatments: control (ambient pollination), reduced pollination, and increased (supplemented) pollination) (Kearns & Inouye 1993). In the reduced treatment, we placed mesh bags over half of flower buds on each plant to exclude pollinators (rounding up for odd numbers). In the increased treatment, we left flowers open for natural pollination and also hand pollinated all receptive flowers three times per week. Plants in the control treatment were not manipulated. For 4–5 years, we censused survival, plant size, flowering status, and seed production of every individual in each treatment (at least 250 individuals per species × treatment). We also censused and tagged seedlings as they emerged. *Hydrophyllum* was treated and censused for five years, the other species for four years. To estimate recruitment rate, we sowed known quantities of seed in separate, adjacent plots and recorded recruitment. To estimate seed bank survival, we buried seed bags and assessed germination rates after 1 to 3 years. Only *Hydrophyllum* appeared to have a seed bank, and this was incorporated into our population model for that species.

### Demographic models

We built a demographic model tailored to each species’ life cycle in two steps: modeling vital rates as a function of plant size and treatment, then using these vital rates to parameterize an Integral Projection Model (IPM). First, to estimate treatment- and size-specific vital rates, we used hierarchical, multivariate linear models. These model vital rates (e.g., survival, flowering probability) as a multivariate response, using treatment as a predictor and size as a covariate. We fit these models in a Bayesian framework using package *brms* (v.2.22.0, Bürkner 2017) to interface between R (v. 4.5.1, R Core Team 2025) and Stan (v.2.36, Gelman *et al*. 2015) (see Supplement for model details).

Next, we composed the demographic vital rate functions into an IPM projection kernel for each species and parameterized it with 1000 joint posterior draws from the fitted vital rate model using the R package *ipmr* (v.0.0.7; Levin *et al*. 2021) (see Supplement and Iler et al. 20XX for details). We projected each of the resulting kernels to obtain posterior distributions of λ and each life history metric (see following section). The Bayesian approach enabled us to propagate uncertainty from the individual-level field census to the vital rate functions, IPM kernels, and demographic statistics.

### Life history metrics

To assess the effects of pollination treatment on life histories, we estimated metrics related to life history strategy (Salguero-Gómez *et al*. 2016) and demographic resilience (Capdevila *et al*. 2020, 2022). Following Salguero-Gómez et al.’s (2016) framework for summarizing life history strategy, we estimated eight life history metrics related to the slow-fast continuum and “reproductive strategy” (Table 1). Following Capdevila et al.’s (2020) framework for quantifying demographic resilience, we estimated three life history metrics to describe demographic compensation, resistance, and rate of return to equilibrium (Table 1).

**Table 1.**
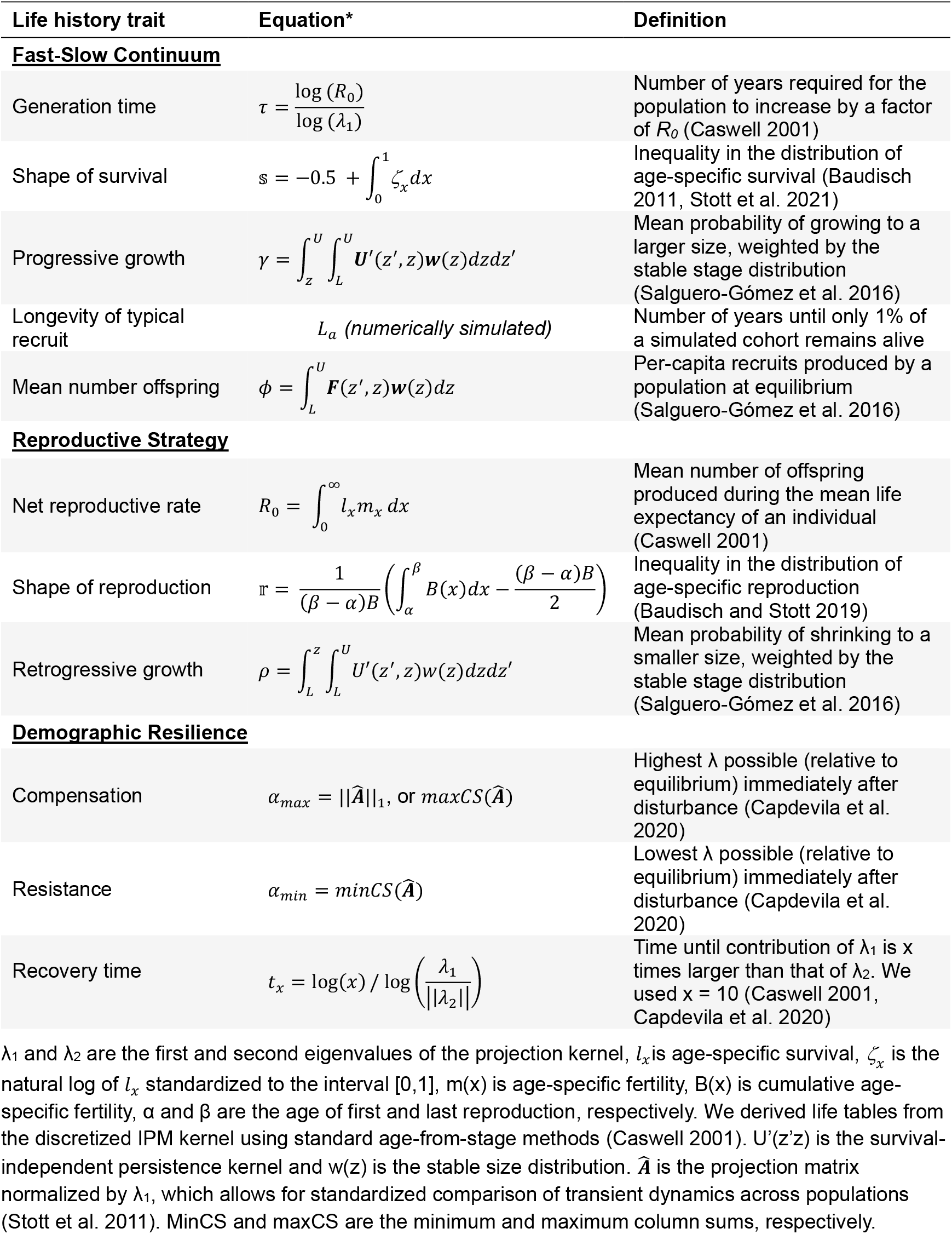
Life history traits measured for each experimental population.

In total, we calculated 11 life history metrics (Table 1) and the population growth rate, λ. We refer to these all together as ‘demographic statistics.’ To estimate posterior distributions for each demographic statistic in each treatment, we calculated the statistic for each posterior draw of the IPM kernels using the R packages Rage (v.1.8.0, Jones *et al*. 2022) and popdemo (v.1.3-2, Stott *et al*. 2021). For metrics that require a starting size, we used the posterior mean recruit size.

### Effects of pollination treatment

To characterize the multivariate life history response of each plant species to the pollination treatments, we used canonical discriminant analysis (CDA). Briefly, CDA finds a linear combination of variables that maximizes separation among groups, similar to a multivariate ANOVA. In this case, our groups are the treatments, our variables used to separate treatments are the demographic statistics, and our data are the samples (draws) from the joint posteriors of those statistics. In practical terms, the CDA summarizes how the 12 demographic statistics differ among treatments. To run the CDA, we first run a multivariate regression using the function lm, then rotate the data and calculate variable loadings using the package *candisc* (v. 0.9.0, Friendly 2007; Friendly & Fox 2024). We visualized the CDA using ordination biplots. We used these biplots to visualize the differences among treatments, to ask which demographic statistics contribute most to those changes, and which of these were consistent across species.

Additionally, to assess changes in individual demographic statistics, we examined marginal posterior contrasts between each manipulated treatment and the control. That is, the posterior distribution for the difference between the treatment and the control. In the Supplemental Material, we also report direct treatment contrasts—the difference between the reduced and increased treatments. This contrast provides a clearer comparison of life history under low versus high pollination but does not speak as directly to *change* from current conditions. In both cases, we measure the strength of evidence for each difference using posterior credible intervals: we considered there to be strong evidence for a treatment effect if the 90% credible interval of the difference did not include 0, and we considered there to be weak evidence for a treatment effect if the 50% credible interval did not include 0. We were hesitant to draw conclusions based on weak evidence for one change by itself (e.g., if there is weak evidence for a change in only one species), but we did interpret weak evidence for the same response across species as evidence for a trend (e.g., if all four species show weak evidence for the same response). We also give such results more weight if reduced versus increased pollination treatments show opposing trends.

## Results

Changing pollination service caused shifts in life history strategy in all four plant species. This is evident from the divergence of joint posterior distributions of our demographic statistics (Figure 1) and treatment contrasts of individual life history traits (Figures 2, S1-2). Pollination treatment responses were widespread among species, with each species showing evidence of change in 2–9 (of 11) life history metrics. Likewise, life history was broadly impacted by pollination treatments, with all 11 metrics we measured changing in at least one species.

**Figure 1.**
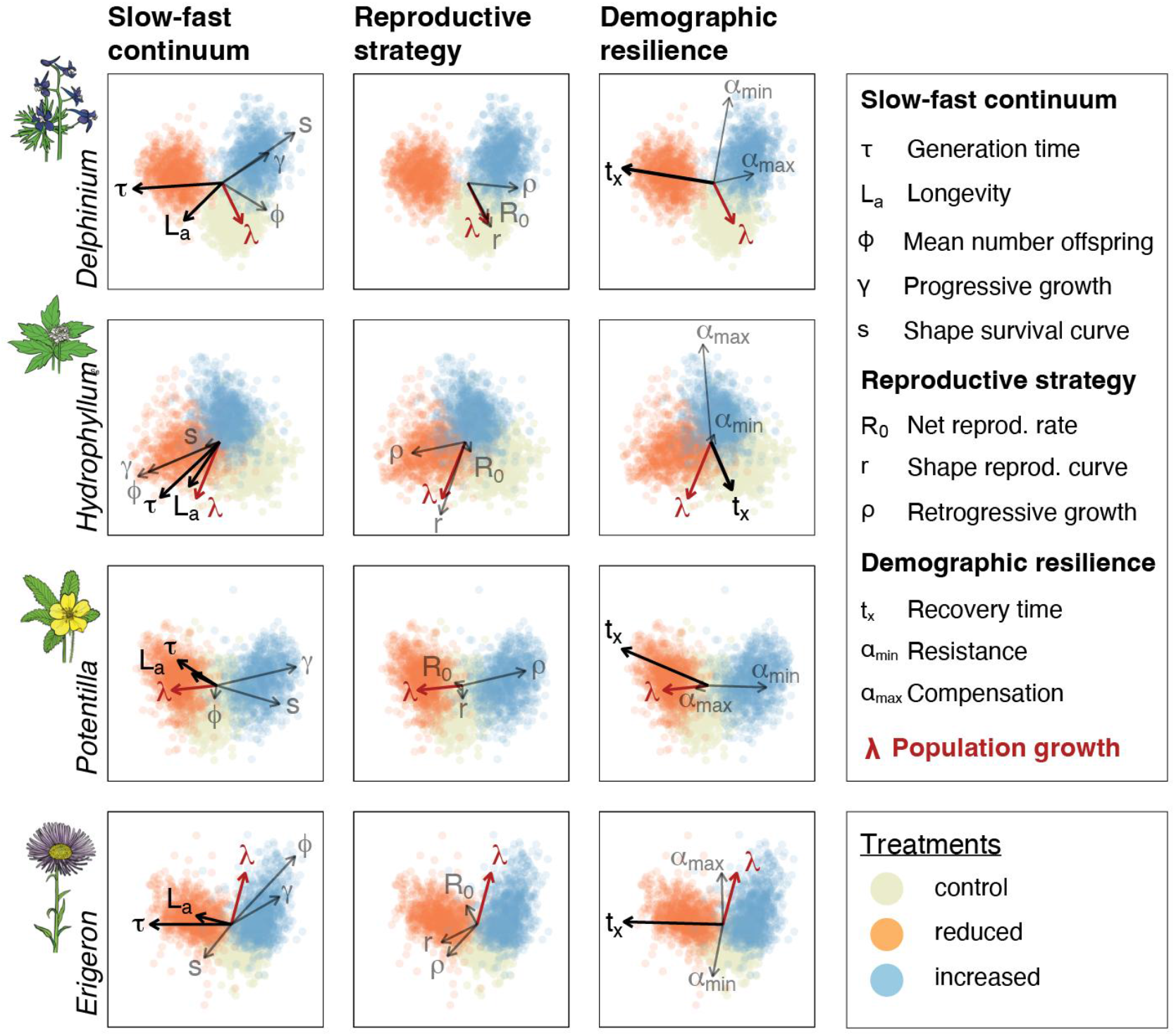
Canonical discriminant analysis of the joint posterior distributions of our 12 demographic statistics. Colored points are draws from the joint posterior for the 12 statistics, and the arrows are loadings for each statistic. The direction and magnitude of each arrow indicate the how that statistic changed among treatments (see Figure 1). To improve readability, the biplot for each species (rows) is split into three panels (columns): each column displays a subset of loadings for the three categories of life history traits. The black, bolded arrows highlight the three metrics that changed most consistently in response to treatment: generation time, longevity and recovery time. Note: a small number (≤ 1%) of outlying posterior scores were omitted to reduce white space. Plant illustrations by Life Science Studio (J. Johnson).

**Figure 2.**
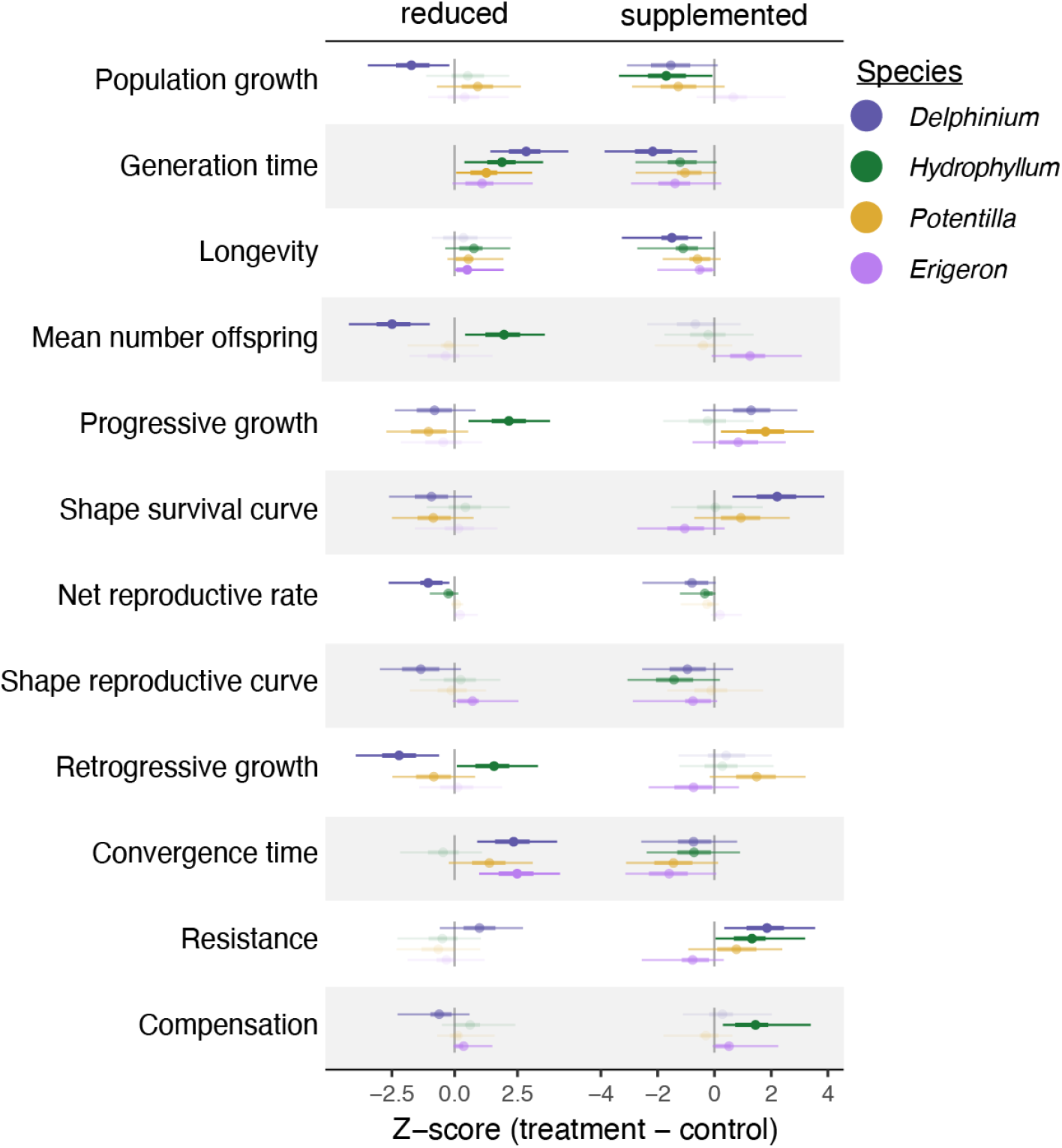
Effects of pollination treatments (treatment – control) on life history traits of our focal plant species. Points represent the posterior mean and thick and thin lines represent the 50% and 90% posterior credible intervals. Faded symbols represent those where the 50% CI overlaps 0 (no evidence of a difference), semi-transparent symbols are those where the 50% CI does not overlap 0 (weak evidence) and the bold symbols are those for which the 90% CI does not overlap 0 (strong evidence). See Table 1 for the definition and calculation of each life history trait. Scaled life history trait values for all treatment levels are shown in Fig. S1 and raw values are summarized in Table S2.

The most consistent effects of pollination treatment were on generation time (τ), longevity (*L*_*α*_) and recovery time (*t*_*x*_). High and low pollination treatments were consistently distinguished by these three traits (bolded arrows in Figure 1; direct treatment contrasts in Figure S2), with the exception that recovery time was not associated with pollination treatment in *Hydrophyllum*. Generation time, longevity, and recovery time were all greater in the reduced pollination treatment than in the increased pollination treatment. Contrasting each treatment to the control (Figure 2), we see the same result, with the exception that *Delphinium* longevity did not differ between the reduced treatment and the control.

The magnitude of these life history responses was biologically meaningful. Reducing pollination increased generation time by at least 19% (6.1 years, for *Hydrophyllum*) and up to 44% (8.2 years, for *Erigeron*) relative to the control. The impact of increased pollination on generation time was weaker, but still decreased by at least 10% (–3.8 years, for *Hydrophyllum*) and as much as 23% (–4.5 years, for *Erigeron*). Similarly, longevity (*L*_*α*_) increased between 16% (4.2 years, for *Hydrophyllum*) and 140% (450 years, for *Erigeron)* in the reduced treatment and decreased between 18% (–8.0 years, for *Potentilla*) and 38% (–130 years, for *Erigeron*) in the increased treatment, and recovery time increased by 24% (0.51 years, for *Potentilla*) to 78% (1.5 years, for *Delphinium*) in the reduced treatment and decreased by 11% (0.25 years, for *Delphinium*) to 20% (0.47 years, for *Potentilla*) in the increased treatment. See Table S2 for a complete summary of demographic statistics under each treatment.

Other changes in life history were inconsistent among species (Figures 1 and 2). For example, under reduced pollination, retrogressive growth ( *ρ*) decreased for *Delphinium* and *Potentilla* but increased for *Hydrophyllum*. Similarly, under increased pollination, demographic resistance (α_min_) decreased for *Erigeron* but increased for the other three species. Notably, changes in population growth rate (λ) were also inconsistent across species (see also: Iler et al 20XX).

## Discussion

Environmental changes can directly impact multiple aspects of a species’ life history strategy. Here, we found changes in pollination rate to affect nearly every aspect of plant life history. Although many effects were idiosyncratic across species – including effects on population growth rate (λ) – reduced pollination rate consistently increased longevity, generation time and recovery time, while increased pollination did the opposite.

The impact of pollination change on plant life history was magnified by a tradeoff between reproduction versus growth and survival. By manipulating the amount of pollen received by plants, we altered seed production—essentially coercing a change in plants’ reproductive investment. And, because the resources available to the plants did not change, this change in reproductive investment traded off with investment in interannual growth and/or survival (Figure S3; Iler et al 20XX). This tradeoff had two main consequences. First, changes in pollination did not lead to consistent changes in λ (Figures 1, 2, S2; Iler et al 20XX). Second, changes in survival and reproductive schedules led to changes in emergent life history traits, including consistent shifts in generation time, longevity, and recovery time.

Notably, these changes in life history among pollination treatments resemble differences in life history among species. Among species, life history variation is largely structured by the tradeoff between reproduction versus survival and growth (or current vs. future reproduction; Stearns 1976; Van Noordwijk & De Jong 1986; Williams 1966). This tradeoff leads to the slow-fast continuum of life histories, with slow species characterized by – among other things – long lives and generation times, and fast species characterized by short lives and generation times (Gaillard *et al*. 2005; Healy *et al*. 2019; Pianka 1970; Salguero-Gómez *et al*. 2016). These are precisely the changes that were most consistent among our pollination treatments: reduced pollination led to increased longevity and generation time (i.e., slower pace of life), and increased pollination led to decreased longevity and generation times (i.e., faster pace of life).

The observed shifts in generation time and longevity are likely to have their own downstream consequences. For example, we expect the slower-living populations under reduced pollination to be less sensitive to environmental stochasticity (Stearns 1976), but also be slower to recover from disturbance or even adapt to novel conditions (Cotto *et al*. 2017; Kuparinen *et al*. 2010; Nordstrom & Melbourne 2025; Yue *et al*. 2010). The opposite is also true: the faster-living populations under increased pollination would be more sensitive to environmental stochasticity but also be faster to recover and adapt. These changes in life history thus represent not just changes in how these plants live their lives, but also how their populations would relate to their environment moving forward. This is especially relevant now, given that climate change is expected to result in increasingly novel and variable environmental conditions (Calvin *et al*. 2023; Williams & Jackson 2007).

All that said, distilling the effects of pollination on life history down to only “slow” versus “fast” would be overly simplistic. Other than generation time and longevity, traits typically associated with the slow-fast continuum changed inconsistently across species. We also analyzed traits describing “reproductive strategy” and demographic resilience (Capdevila *et al*. 2020; Salguero-Gómez *et al*. 2016). All these metrics changed in some way but, again, these changes were inconsistent across species. For example, in comparing the reduced to increased pollination treatments, the rate of retrogressive growth increased with higher pollination in two species (*Delphinium* and *Potentilla*), decreased with higher pollination in a third (*Hydrophyllum*), and did not seem to change in the fourth (*Erigeron*) (Figures 1, S2). Similarly, demographic resistance, which describes a population’s immediate sensitivity to disturbance, increased with increasing pollination in three species but decreased with increasing pollination in the fourth. Such idiosyncratic changes should perhaps be unsurprising. Life history traits are complex, emergent properties of multiple underlying vital rates, *and* how these vital rates change with plant size. Take the examples of retrogressive growth: In *Delphinium* and *Potentilla*, the effect pollination treatment manifests (in part) as a tradeoff between seed production and size-dependent growth, such that larger plants—those that produce the most seeds—have a lower growth rate under increased pollination and are thus more likely to shrink. Given that most individuals are intermediate to large, this change in growth rate increases the population-level frequency of retrogressive growth. In *Hydrophyllum*, by contrast, pollination treatment affected survival more than growth rate and retrogressive growth decreased. Here, the decrease in retrogressive growth is likely a result of a change in the stable size distribution. Under increased pollination, there become a greater proportion of individuals in the seed bank, who can only grow, not shrink, thereby reducing the population-level frequency of retrogressive growth. More broadly, consider that the responses of individual vital rates to environmental change, and the correlations or tradeoffs among them, vary among species (Fay *et al*. 2022; Pagel *et al*. 2020; Wilson & Martin 2012) and contexts (Sletvold & Ågren 2015; Villellas & García 2018). As a result, many life history changes might be difficult to predict. And thus, while the effects of pollination on generation time and longevity do resemble shifts along the slow-fast continuum, the remaining traits remind us that life history is complex, and species will respond to environmental change in their own, often unpredictable, ways.

The life history changes we documented here are almost purely plastic. Although mortality and recruitment did occur, individual turnover was limited: 41–83% individuals from the first census were still alive in the final census, and 13–57% of individuals in the final census were not present in the first. Long term, however, these plastic changes might be further augmented by adaptation. Local adaptation in life history has been documented previously (DeMarche *et al*. 2016; Lind *et al*. 2011; Wang *et al*. 2018), including on floral traits (Newman *et al*. 2015; Sletvold *et al*. 2017) and reproductive investment by plants (Frei *et al*. 2014; Hautier *et al*. 2009; Helsen *et al*. 2020). We might expect something similar here, especially in cases where a change in pollination negatively affects λ. For example, three of our species showed evidence for reduced λ in response to increased pollination. If high-pollination conditions continued long term, one possibility for these species is selection for reduced floral investment to offset the cost of reproduction. The life history consequence of such adaptation is unclear, but we expect new conditions would ultimately lead to a new life history optimum (Cayuela *et al*. 2022; Felmy *et al*. 2022; Williams *et al*. 2015). In response to reduced pollination, one species showed evidence for reduced λ. In this case, one possibility is selection for increased floral investment to make flowers more competitive, thereby increasing rate but also cost of reproduction. Here too, it is hard to know what effect this might have on life history. Regardless, whether by plasticity or adaptation, the takeaway remains: changes in pollination services are likely to affect plant life history strategy and, in turn, long-term population dynamics.

Understanding population responses to environmental change is an increasingly urgent need. For most plant species, a key aspect of the environment is pollination services by animals. Given the risk of broad-scale pollinator declines (Cornelisse *et al*. 2025), a natural question has been whether pollination services affect λ (Baer & Maron 2018; Knight 2004; Law *et al*. 2010). Yet changing λ might not be the only – or the most consistent – response to changing pollination. Here, we found changes in pollination rates affect plant life history more consistently than they affect λ. In all four of our study species, changing pollination services led to shifts in generation time and longevity that resemble shifts along the slow-fast continuum, in addition to other pervasive but idiosyncratic changes. By inducing a life history tradeoff, altered pollination services effected change within populations that resembled life history differences among species. Our results highlight life history as an important but underappreciated dimension to the effects of environmental change.

## Acknowledgements

We thank the many field assistants and students who helped collect data in the field, the Iler-CaraDonna lab group for feedback on earlier versions of the manuscript, and the CBG Train Club for fruitful discussions of life history. This work was supported by NSF DEB 1754518 to AMI and PJC, NSF DEB 2211721 to AMI, and the Research Capacity Fund (HATCH) project award no. 7004646 to WKP from the U.S. Department of Agriculture’s National Institute of Food and Agriculture.

λ_1_ and λ_2_ are the first and second eigenvalues of the projection kernel, *l*_*x*_ is age-specific survival, *ζ*_*x*_ is the natural log of *l*_*x*_ standardized to the interval [0,1], m(x) is age-specific fertility, B(x) is cumulative age-specific fertility, α and β are the age of first and last reproduction, respectively. We derived life tables from the discretized IPM kernel using standard age-from-stage methods (Caswell 2001). U’(z’z) is the survival-independent persistence kernel and w(z) is the stable size distribution. **Â**is the projection matrix normalized by λ_1_, which allows for standardized comparison of transient dynamics across populations (Stott et al. 2011). MinCS and maxCS are the minimum and maximum column sums, respectively.

## Supplement

### Modeling methods

We built our population models using a combination of multivariate linear models and integral projection models (IPMs). First, we modeled vital rates – survival from time t to t+1, growth (size at time t+1), the probability of flowering, and number of seeds produced by flowering plants – as a function of plant size at time t and treatment. We also modeled seedling recruitment probability and seedling size as a function of treatment. These models were run together as a single multivariate model using Stan and the R package *brms*. Because most sub-models were generalized models, using binomial or negative binomial error distributions, we could not estimate correlations among model residuals. Nevertheless, this approach still has the benefit of retaining an inherent link in the joint posterior probability distribution across parameters and treatments. We ran models on four chains for 2000 iterations warm up and 2000 iterations sampling, resulting in 8000 posterior draws with which to characterize the posterior distribution of each parameter. We assessed convergence visually and by assuring all R-hat < 1.05.

Second, we used the results of our vital rate models to parameterize integral projection models (IPMs). We estimated posterior distributions of IPM kernels and derived parameters by parameterizing 1000 IPMs using independent draws from the joint vital rate posterior distribution. We discretized these kernels using 400 mesh points spread evenly across the observed size bounds extended by 15%, yielding a 400 x 400 matrix projection model that described plants 15% larger and smaller than any we observed. We corrected for eviction by truncating the distribution of *t+1* size at the expanded size bounds.

^a^(Waser and Price 1990) ^b^(Ingold et al. 2024) ^c^(Beckmann Jr. 1979) ^d^(Vail 1983) ^e^(Iler, unpub. data) ^f^(Iler et al. in prep)

**Table S1.** Reproductive characteristics of the five perennial study species.

| Species | Self-compatible | Autogamous selfing | Locally pollen limited | Seed bank |
| --- | --- | --- | --- | --- |
| *Delphinium nuttallianum*<br>(Ranunculaceae) | yes<sup>a</sup> | no<sup>a</sup> | yes<sup>a</sup> | no<sup>f</sup> |
| *Erigeron speciosus*<br>(Asteraceae) | no<sup>b</sup> | no<sup>b</sup> | no<sup>b</sup> | no<sup>f</sup> |
| *Hydrophyllum fendleri*<br>(Boraginaceae) | yes<sup>c</sup> | no<sup>c</sup> | yes<sup>e</sup> | yes<sup>f</sup> |
| *Potentilla pulcherrima*<br>(Rosaceae) | yes<sup>d</sup> | yes<sup>d</sup> | yes<sup>e</sup> | no<sup>f</sup> |

<sup>a</sup>(Waser and Price 1990) <sup>b</sup>(Ingold et al. 2024) <sup>c</sup>(Beckmann Jr. 1979) <sup>d</sup>(Vail 1983) <sup>e</sup>(Iler, unpub. data) <sup>f</sup>(Iler et al. in prep)

**Figure S1.**
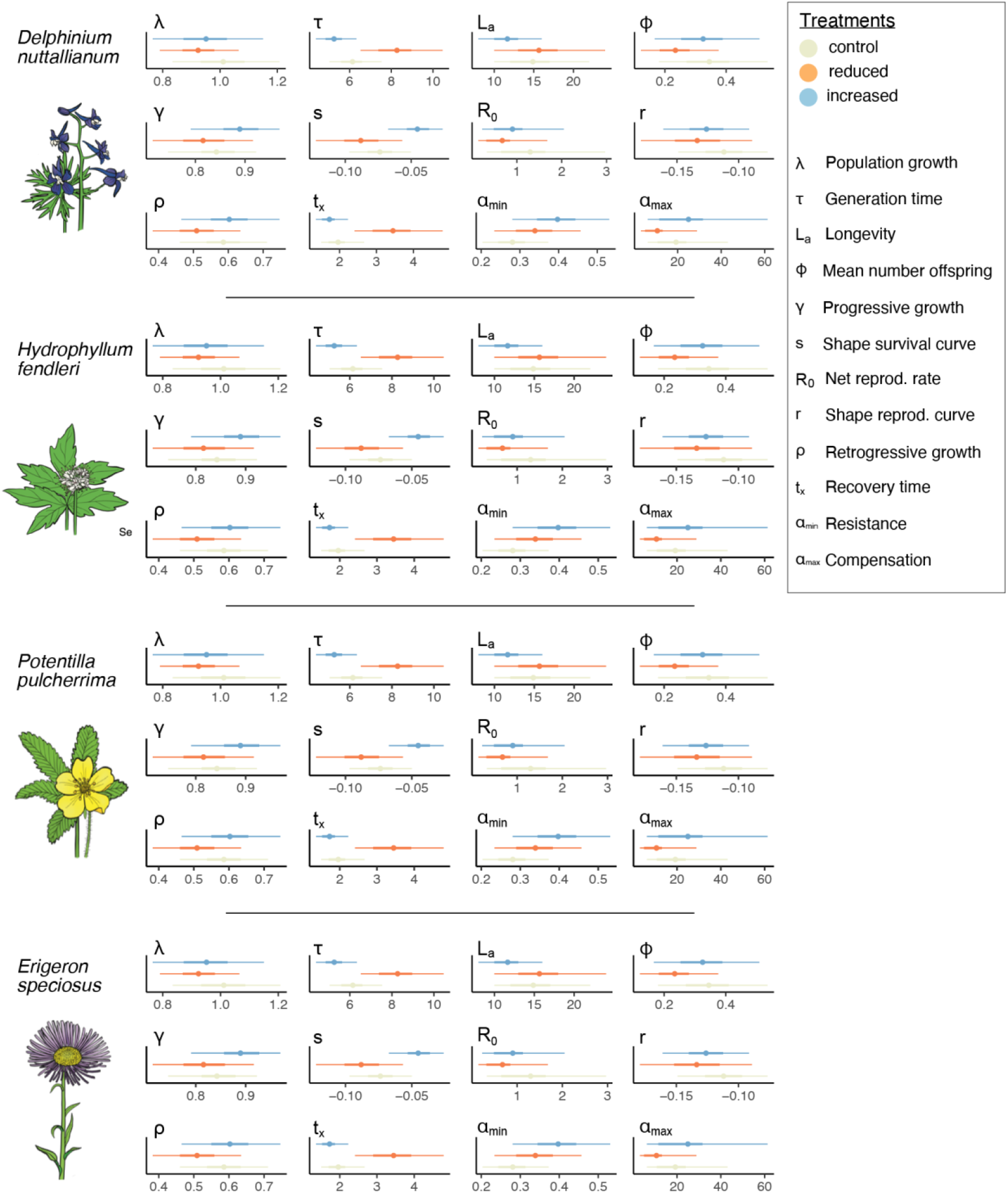
Marginal posteriors for our 12 demographic statistics. Points represent mean values and thick and thin bars represent 50% and 90% credible intervals. Note: marginal posteriors should not be used to compare treatments, because individual posterior draws are linked. To compare treatments, use the treatment contrasts in Figures 3 and S2. Plant illustrations by Life Science Studio (J. Johnson).

**Figure S2.**
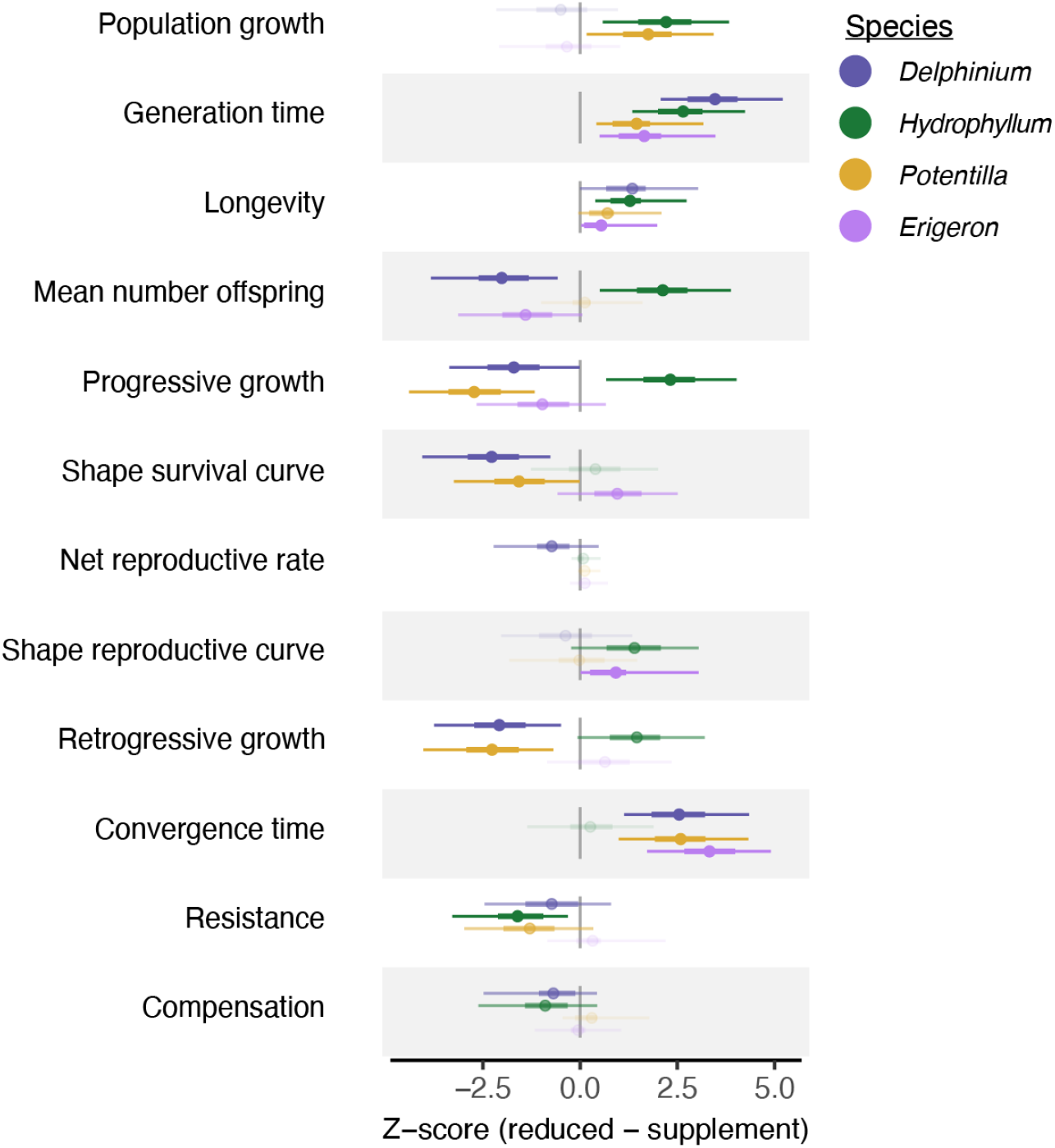
Direct treatment contrasts for each life history trait and species. Values greater than 0 mean the trait value was higher in the reduced treatment; values lower than 0 mean the trait value was higher in the supplemented treatment. Points represent the posterior mean and thick and thin lines represent the 50% and 90% posterior credible intervals. Faded symbols represent those where the 50% CI overlaps 0 (no evidence of a difference), semi-transparent symbols are those where the 50% CI does not overlap 0 (weak evidence) and the bold symbols are those for which the 90% CI does not overlap 0 (strong evidence). See Table 1 for the definition and calculation of each life history trait. Scaled life history trait values for all treatment levels are shown in Fig. S1 and raw values are summarized in Table S2.

**Figure S3.**
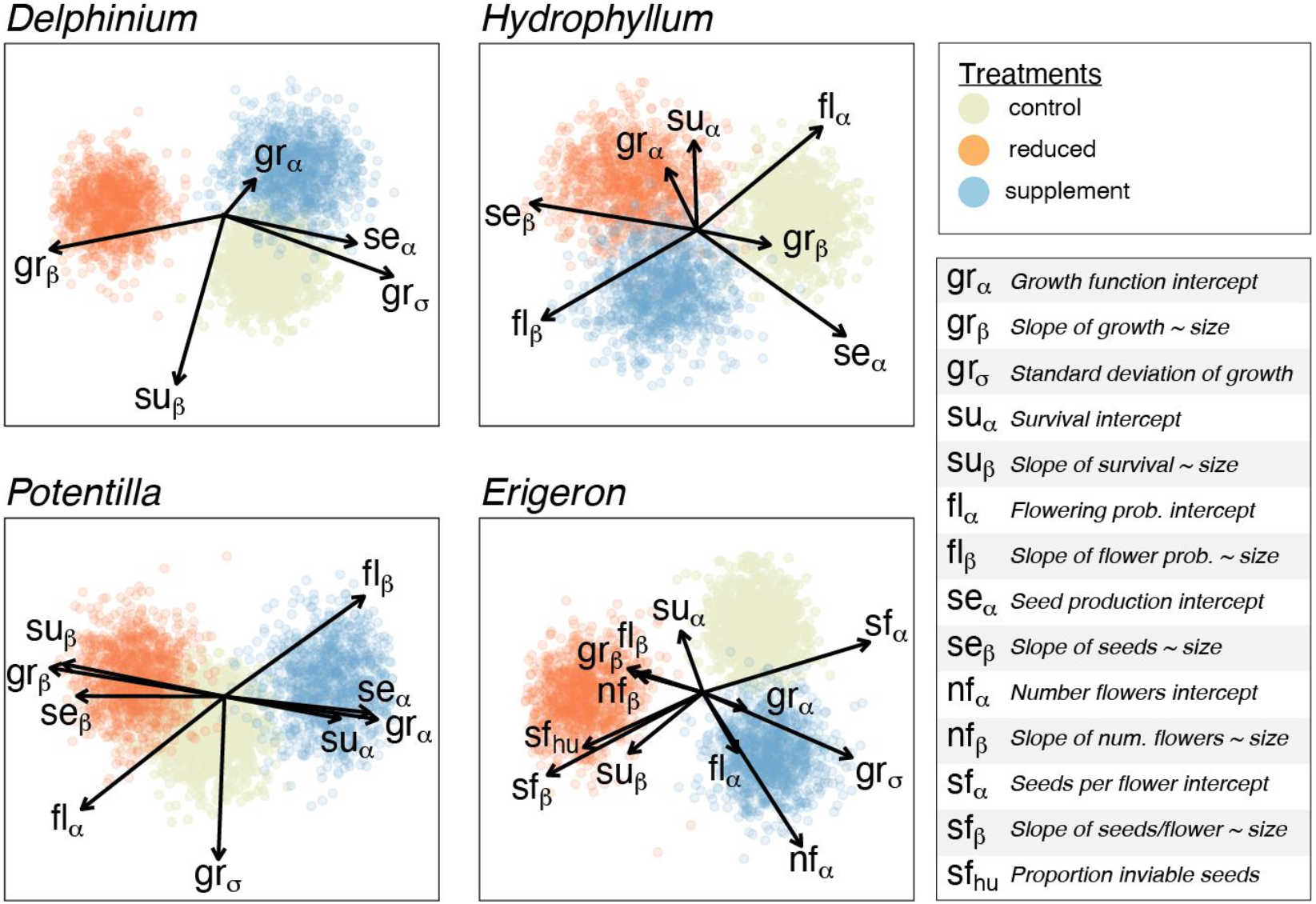
Canonical discriminant analysis of treatment separation by vital rate model parameters. Models for all species included the vital rates growth (size in t+1 ∼ size in t) and survival from t to t+1. Variance around average growth is explicitly included. For *Delphinium, Hydrophyllum*, and *Potentilla*, the flowering model included probability of flowering and seeds produced conditioned on a the plant flowering. For *Erigeron*, which is in *Asteraceae* and has composite flowers, the flowering model included the number of heads per plant, seeds per seeds per head, and the proportion of failed or inviable seeds. Every vital rate (except proportion failed seeds) was modeled as a function of plant size and so includes an intercept (α) and slope (β). Variance around average growth is also included (*gr*_σ_). Not all panels include all parameters because loadings that did not contribute to the difference among treatments (and were thus clustered tightly in the center) were removed to improve legibility. Notice the tradeoffs between seed production and growth: In *Delphinium*, baseline seed production trades off between treatments with the slope on growth. A higher slope on growth means greater growth for mid-sized plants and reduced shrinkage for large plants – essentially, greater somatic investment by plants large enough to flower. In *Hydrophyllum*, baseline growth and survival trade off with seed production. In *Potentilla*, baseline seed production trades off with the slopes on growth and survival. In *Erigeron*, the number of flowers and seeds per fruit trade off with baseline survival and the slopes on survival and growth.

## Notes

### Competing Interest Statement

The authors have declared no competing interest.

